# Unifying approaches from statistical genetics and phylogenetics for mapping phenotypes in structured populations

**DOI:** 10.1101/2024.02.10.579721

**Authors:** Joshua G. Schraiber, Michael D. Edge, Matt Pennell

## Abstract

In both statistical genetics and phylogenetics, a major goal is to identify correlations between genetic loci or other aspects of the phenotype or environment and a focal trait. In these two fields, there are sophisticated but disparate statistical traditions aimed at these tasks. The disconnect between their respective approaches is becoming untenable as questions in medicine, conservation biology, and evolutionary biology increasingly rely on integrating data from within and among species, and once-clear conceptual divisions are becoming increasingly blurred. To help bridge this divide, we derive a general model describing the covariance between the genetic contributions to the quantitative phenotypes of different individuals. Taking this approach shows that standard models in both statistical genetics (e.g., Genome-Wide Association Studies; GWAS) and phylogenetic comparative biology (e.g., phylogenetic regression) can be interpreted as special cases of this more general quantitative-genetic model. The fact that these models share the same core architecture means that we can build a unified understanding of the strengths and limitations of different methods for controlling for genetic structure when testing for associations. We develop intuition for why and when spurious correlations may occur using analytical theory and conduct population-genetic and phylogenetic simulations of quantitative traits. The structural similarity of problems in statistical genetics and phylogenetics enables us to take methodological advances from one field and apply them in the other. We demonstrate this by showing how a standard GWAS technique—including both the genetic relatedness matrix (GRM) as well as its leading eigenvectors, corresponding to the principal components of the genotype matrix, in a regression model—can mitigate spurious correlations in phylogenetic analyses. As a case study of this, we re-examine an analysis testing for co-evolution of expression levels between genes across a fungal phylogeny, and show that including covariance matrix eigenvectors as covariates decreases the false positive rate while simultaneously increasing the true positive rate. More generally, this work provides a foundation for more integrative approaches for understanding the genetic architecture of phenotypes and how evolutionary processes shape it.

## Introduction

Statistical genetics and phylogenetic comparative biology share the goal of identifying correlations between features of individuals (or populations) that are structured by historical patterns of ancestry. In the case of statistical genetics, researchers search for causal genetic variants underlying a phenotype of interest, whereas in phylogenetic comparative biology, researchers are typically interested in testing for associations among phenotypes or between a phenotype and an environmental variable. In both cases, these tests are designed to isolate the influence of a focal variable from that of many potential confounding variables. But despite the shared high-level goal, the statistical traditions in these two fields have developed largely separately, and—at least superficially—do not resemble each other. Moreover, researchers in these two statistical traditions may have different understandings of the nature of the problems they are trying to solve.

In statistical genetics, phenotypes and genotypes can be spuriously associated because of confounding due to population structure [1–4] or assortative mating [5, 6]. For example, in their famous “chopsticks” thought experiment, Lander & Schork [1] imagined that genetic variants that have drifted to higher frequency in subpopulations in which chopsticks are frequently used will appear, in a broad sample, to be associated with individual ability to use chopsticks, even though the association is due to cultural confounding and not to genetic causation. Confounding can also be genetic [7]—if a genetic variant that changes a phenotype is more common in one population than others, leading to differences in average phenotype among populations, then other, non-causal variants that have drifted to relatively high frequency in this population may appear to be associated with the phenotype in a broad sample. In addition to affecting genome-wide association study (GWAS) results, such confounding can affect heritability estimation [8, 9], genetic correlation estimates [10, 11], and prediction of phenotypes from polygenic scores [12–16]. Although many candidate solutions have been offered [17–21], the two most common approaches involve adjusting for shared ancestry using the genetic relatedness matrix (GRM, [22]), either by incorporating individual values on the first several eigenvectors of this matrix (i.e. the principal components of the genotype matrix) as fixed effects [23], or by modeling covariance among individuals attributable to genome-wide relatedness in a linear mixed model (LMM, [24–28]).

In phylogenetic comparative biology, researchers typically aim to control for the similarity of related species by incorporating the species tree into the analysis. There has been a great deal of controversy over the years as to what the underlying goals and implicit assumptions of phylogenetic comparative methods (PCMs) are [see for examples refs. 29–36]. But broadly speaking, it seems that many researchers understand the goal of PCMs to be avoiding “phylogenetic pseudoreplication” [37]—mistaking similarity due to shared phylogenetic history for similarity due to independent evolutionary events [34]. This is most commonly done by conducting a standard regression, using either generalized least squares (GLS) or a generalized linear mixed model (GLMM), but including the expected covariance structure owing to phylogeny [38–42]. (Throughout this paper, we do not make a distinction between phylogenetic GLS and phylogenetic GLMM models. We refer to them generically by the shorthand GLS for the general case and PGLS for cases where the phylogenetic variance-covariance is used; see below for definitions.) This covariance structure reflects both the relatedness of species and the expected distribution of phenotypes under a model of phenotypic evolution [43, 44], such as a Brownian motion [45] and related alternatives [44]. (The “phylogenetically independent contrasts” method [46], which ushered in modern PCMs, is statistically equivalent to a PGLS model assuming a Brownian model [47].)

In recent years, however, signs have emerged that these two subfields may benefit from closer conversation, as emerging approaches in both statistical genetics and phylogenetics encounter questions that call for the other subfield’s expertise. For example, in humans, evolutionary conserved sequences are enriched for trait and disease heritability [48, 49], and conservation across related species can be used to prioritize medically relevant variants in fine mapping [50, 51] and rare-variant association studies [52, 53]. Similarly, multi-species alignments are being used by conservation geneticists to estimate the fitness effects of mutations in wild populations [54, 55] and by plant breeders to aid in genomic selection [56, 57]. And there is growing interest in using estimated ancestral recombination graphs (ARGs) to perform explicitly tree-based versions of QTL mapping and complex trait analysis [58, 59]. From the phylogenetics side, researchers are increasingly employing GWAS-like approaches (“PhyloG2P” methods; [60]) for mapping phenotypes of interest for which the variation primarily segregates among rather than within species. Futhermore, phylogenetic biologists have been developing phylogenetic models that consider within-species polymorphisms in genes [61, 62] and how ancient polymorphisms that are still segregating among species may confound association tests [63].

Such emerging connections suggest that it would be beneficial to better understand the ways in which statistical genetics and phylogenetic comparative biology relate to each other. To this end, we start from first principles and develop a general statistical model for investigating associations between focal variables while controlling for shared ancestry and environment. We show that both standard GWAS and standard phylogenetic regression emerge as special cases of this more general model. We illustrate how this deep connection provides insights into why these two classes of methods can be misled by certain types of unmodeled structure, why solutions in both fields work (and when do they not), and how statistical advances in either domain can be applied straightforwardly to the other.

## Results

### A general polygenic model of quantitative trait variation

We assume a standard model in which many genetic factors of small effect influence a phenotype in an additive way—that is, there is no dominance or interaction among genetic loci (epistasis). Denoting by *β*_*l*_ the additive effect size of the variant at the *l*th locus and *G*_*il*_ the genotype of the *i*th individual at the *l*th locus, we write

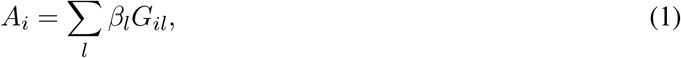

where *A*_*i*_ is the genetic component of the phenotype of individual *i*, sometimes called a genetic value or breeding value. We then express the phenotype of individual *i, Y*_*i*_, as the sum of the genetic component and an environmental component, *E*_*i*_.

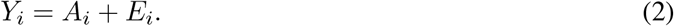

Due to shared ancestry, the genotypes of individuals in the sample will be correlated; thus, the genetic components of the individuals in the sample will be correlated. Moreover, the environments experienced by individuals may be correlated, and these environmental effects may be correlated with the genetic components. If we are interested in understanding the factors that shape the trait of interest, we must control for the covariance induced by shared genetics and shared enviornments. This covariance can be written as

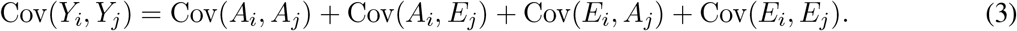

For the rest of the paper, our focus will be on the first term, Cov(*A*_*i*_, *A*_*j*_), the covariance in phenotypes between individuals due to genetic covariance. As we will show below, many models used by both statistical geneticists and phylogenetic biologists can be understood without reference to the components that include environmental effects. Interestingly, some statistical geneticists interpret the use of the GRM in the standard GWAS model as being primarily a proxy for shared (but unmeasured) environment among individuals [7], since in many cases, individuals whose genotypes are correlated will also experience similar environments. Phylogenetic biologists, on the other hand, tend to interpret the use of the phylogenetic covariance matrix in the model exclusively as controlling for genetic effects [29, 33]. On macroevolutionary time scales, it is not so obvious that genetic and environmental similarities will mirror each other. However, at least for traits such as gene expression (which is increasingly studied in a phylogenetic context; [62, 64–67]) that are strongly environmentally dependent and for which measurements are taken from species in different environments, we may need to develop new ways to model the environmental terms. This is a challenge beyond the scope of the present paper. There are some circumstances in which genetic covariance in equation 3 is undefined, such as when effect sizes have an undefined variance [68], or under certain phenomenological models of evolution on phylogenies [69, 70]; we reserve these situations for future work and focus on situations in which the genetic covariance is finite in the subsequent sections.

### Conceptualizations of the genetic covariance

Individuals who are more closely related will have more similar genotypes. For example, individuals in the same local population may share the same alleles identical by descent due to recent common ancestry. On the other hand, individuals in different species may not share alleles due to the species being fixed for alternative alleles at a given locus. Using equation 1,

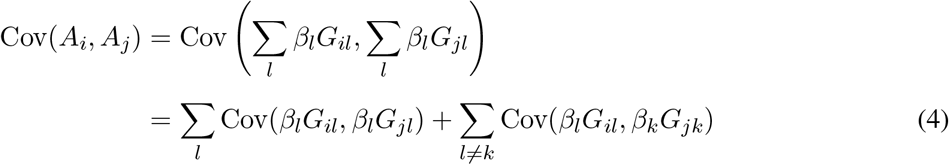

The first term arises from the correlations at a single locus, whereas the second term arises from correlations among loci across individuals. We focus on the first term, though the second can be important in biologically realistic situations. A particularly important case is selection on polygenic traits, which can cause correlations among the genetic contributions to a trait across loci. For example, if a population experiences directional selection on a highly polygenic phenotype, much of the phenotypic change, compared with a related population that has not experienced such selection, is due to to small, coordinated changes in allele frequency [71, 72]. In addition, linkage can affect the evolution of polygenic traits [73] and the results of heritability estimates [74].

We would like to understand how different assumptions about the evolutionary process affect the genetic covariance. However, in full generality, it is hard to say much more, since both the effect sizes and genotypes might be viewed as random and as dependent in arbitrary ways. The particular ways in which effect sizes and genotypes are related is a downstream consequence of assumptions about an underlying evolutionary model. In certain cases, there may be a variable—potentially a latent variable—that renders the genotypes *G*_*il*_ and effect sizes *β*_*l*_ conditionally independent. For example, if the effect size of an allele influences genotypes only through the allele frequency, as is the case under Hardy-Weinberg equilibrium in some models of polygenic selection [75–77], then conditional on the allele frequency, *G*_*il*_ and *β*_*l*_ are independent. We use *Z* to represent such a variable in general. Conditioning on *Z* and using the definition of covariance and the law of total expectation, the first term becomes

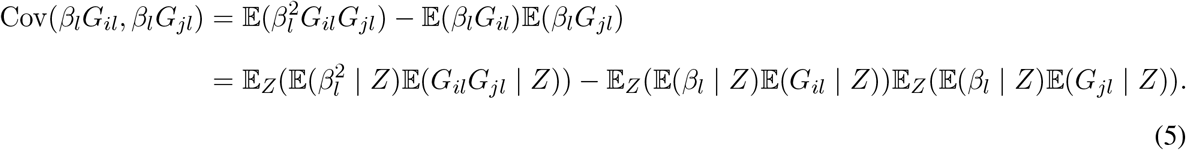

This formula is fully general, as long as the genetic covariance exists, and can be applied in any evolutionary model in which there is a variable *Z* that accounts for the relationship between effect sizes and genotypes. Moreover, it applies when the variable *Z* = *β* or *Z* = *G*.

We will explore how applications across statistical and evolutionary genetics specialize (5) in different ways to create a matrix summarizing genetic covariance in phenotype, which we refer to as Σ. In a sample of *n* individuals (or *n* species), Σ is *n* × *n*, and Σ_*ij*_ is proportional to some version of equation 5. We will see that assumptions made in different fields relate to underlying assumptions about the underlying evolutionary process shaping genetic and phenotypic variation.

Among other names, in different settings, Σ might take the form of a “genetic relatedness matrix,” “kinship matrix,” “expected genetic relatedness matrix,” or “phylogenetic variance-covariance matrix.” Below, we consider the off-diagonal entries of each of these matrices in turn.

### The genetic relatedness matrix

We can simplify equation 5 if the second term is equal to 0, which can be accomplished in a number of ways. One way is to mean-center the genotypes. In that case, we write

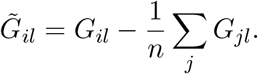

Then, the linear model of the phenotype (equation 2) can be written

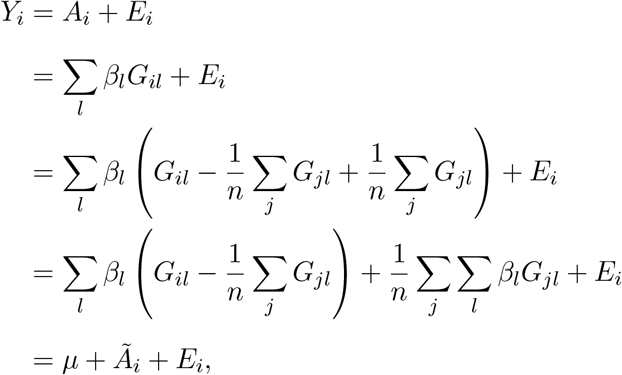

where 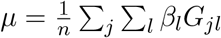 is the mean genetic value in the population, and 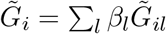. Then, equation 5 becomes

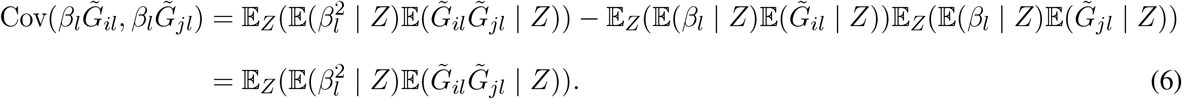

The second line follows because mean-centering implies that 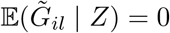 for all *i*.

One reasonable candidate for *Z*—i.e. a variable conditional on which the genotype and effect sizes are independent—is the allele frequency *p*_*l*_. Conditioning on *p*_*l*_ and using mean-centered genotypes,

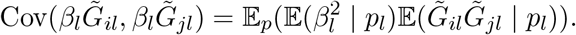

In the presence of dense genotype data, we can approximate the expectation over genotypes by the empirical mean,

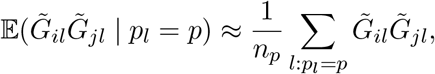

where *n*_*p*_ is the number of sites with frequency *p*. Then, if we have a functional form of 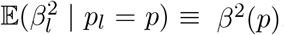, we can compute

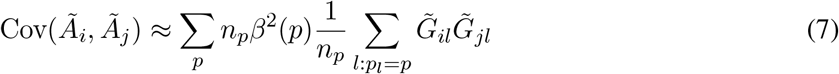

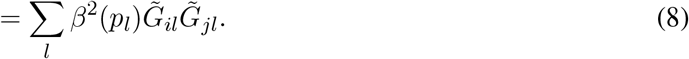

A common choice for *β*^2^(*p*) is

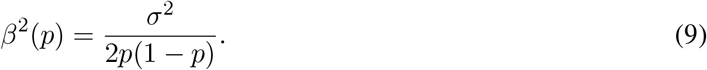

This choice recapitulates the canonical Genetic Relatedness Matrix (GRM), a realization of Σ that is commonly used to estimate heritability from SNP data [78] or to accommodate covariance due to relatedness in GWAS [24–26, 28]. To motivate the relationship in equation 9, in the Appendix we include an argument that the effect size of a locus at frequency *p* under deterministic mutation-selection balance for Gaussian stabilizing selection is

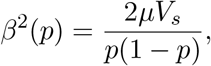

where *μ* is the mutation rate at that locus and *V*_*s*_ is the variance of the Gaussian fitness function. This derivation suggests that equation 9 may be justifiable when the strength of selection is strong enough that genetic drift can be ignored, but may not be appropriate when traits evolve under weaker selection with a substantial contribution of genetic drift [75, 79] or when the model is violated in other ways.

An additional reason for this choice of normalization is that it enables estimation of the additive genetic variance via estimation of *σ*^2^. To see that, note that the additive genetic variance is

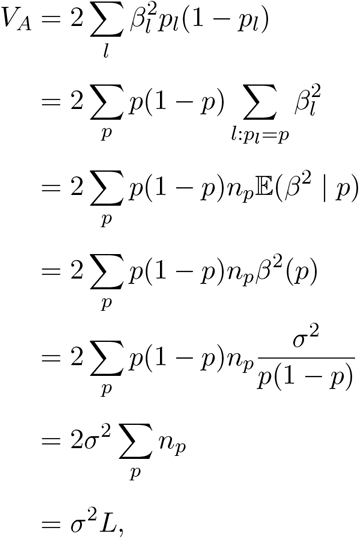

where the second line is arises by grouping variants by allele frequency, the third line comes from recognizing the 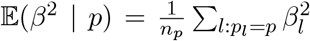 where *n*_*p*_ is the number of loci with frequency *p*, and the final line arises because Σ_*p*_ *n*_*p*_ = *L* is the number of loci. Because of this, approaches that use this normalization of the GRM will tend to overestimate the genetic variance, and hence the heritability, if the true relationship between allele frequency and effect size is weaker than supposed here. This model can be generalized by writing 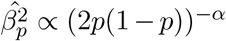, known as the “alpha model” or “LDAK model” [22, 74]. In plant and animal breeding, where this framework first appeared [80], sometimes the same normalization is used as in human genetics, and sometimes genotypes are mean centered but not standardized [81, 82].

### The (pedigree-based) kinship matrix

Historically, plant and animal breeders, along with human and behavior geneticists interested in resemblance of relatives, have frequently faced a situation in which they have had 1) (at least partial) pedigree data describing the parentage of sets of individual plants or animals, 2) phenotypic data on those individuals, but 3) no genome-wide genetic information. In such a situation, one can model the entries of Σ as a function of expected genetic similarity based on the pedigree information, as opposed to realized genetic sharing observed from genotypes [81, 83–85]. One can specialize equation 4 by fixing the effect sizes, leading to

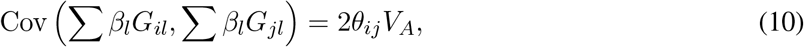

where *θ*_*ij*_ is the kinship coefficient (obtained from the pedigree) relating individuals *i* and *j*, and *V*_*A*_ is the additive genetic variance. Although many derivations exist in standard texts [e.g. 85, 86], we include one in the Appendix for completeness.

Methods based on this formulation include the “animal model” [83, 85, 87, 88], a widely used approach for prediction of breeding values in quantitative genetics. The connection between the animal model and genome-wide marker-based approaches was plain to the quantitative geneticists who first developed marker-based approaches to prediction [80], and it is also noted in papers aimed at human geneticists [22, 78, 89], whose initial interest in the framework focused on heritability estimation. Similarly, the animal model is known to be intimately connected to the phylogenetic methods we discuss later [40–42]. One implication is that close connections between methods used in statistical genetics and phylogenetics, which are our focus here, must exist.

### The expected genetic relatedness matrix (eGRM)

We may make other assumptions to model the genetic covariances among individuals. For instance, we might let *Z* = *β*, the effect sizes themselves. Unlike in the previous subsection, the effect sizes are not fixed; they are random and independent of genotype. In this case, the covariance is computed with respect to alternative realizations of the mutational process (as in the branch-based approach in [90]) and, in this subsection, also alternative realizations of the underlying gene tree(s). In contrast to the assumptions of the previous subsections, independence between the effect sizes and genotypes arises most naturally from a model of neutral evolution.

Then, if genotypes are independent of effect sizes and the expected effect size is 0, we show in the Appendix that

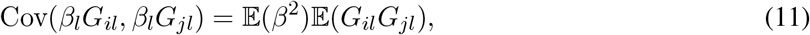

so that, if all loci are equivalent

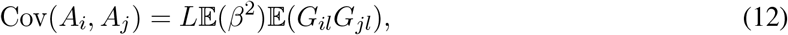

where *L* is again the number of loci. In theory, the expectation of the product of genotypes can be computed based on coalescent theory [91], focusing on the gene trees that describe the pattern of shared inheritance of alleles rather than the distribution of the alleles themselves. In practice, the gene trees underlying genetic variation in the sample cannot be observed directly. However, developing a coalescent approach provides theoretical understanding in its own right, and it also forms a basis for doing complex trait analyses using estimated genome-wide gene trees [58, 59].

Here we develop the gene-tree-based view using a different derivation from that presented by McVean [91]. For simplicity, we assume haploid genetic data, the extension to diploid data is straight-forward but tedious [91]. This argument depends crucially on the assumption that genetic variation is neutral, as in [91]. Incorporating natural selection into gene-tree based models in full generality requires analysis of the ancestral selection graph [92]. First, let 𝒯 be a tree (including branch lengths) and ℚ be the measure on tree space induced by the population history. We use measure-theoretic notation here because trees are a combination of a discrete branching structure and continuous branch lengths; the density *d*ℚ(𝒯) can be roughly thought of as the probability of a tree with a given topology and given branch lengths. To model mutation, we assume that, conditional on the genealogy, mutations arise on each branch as a Poisson process with rate *μ*; thus, given the total branch length in the tree, *T*, the total number of mutations on the tree is distributed a Poisson random variable with mean *μT*. To obtain the density of trees in the infinite-sites limit, we send the mutation rate to 0,

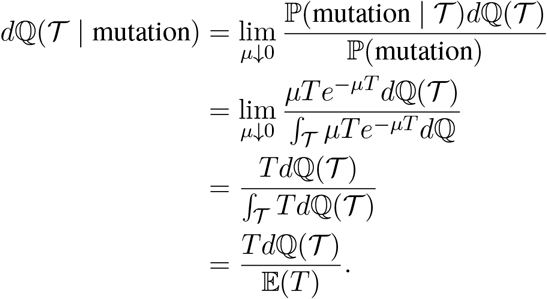

This formula indicates that conditioning on a gene tree having a mutation results in trees with more total branch length. Hence, the distribution of gene trees that underlie SNPs is different from the unconditional distribution of gene trees; in particular, gene trees that have a SNP will tend to have more total branch length than those that do not have a SNP.

Next, note that two haploid individuals will have the same genotype at a variable site only if that mutation occurred in a branch that is ancestral to both samples. Letting *T*_*ij*_ be the time in branches ancestral to both samples, we have

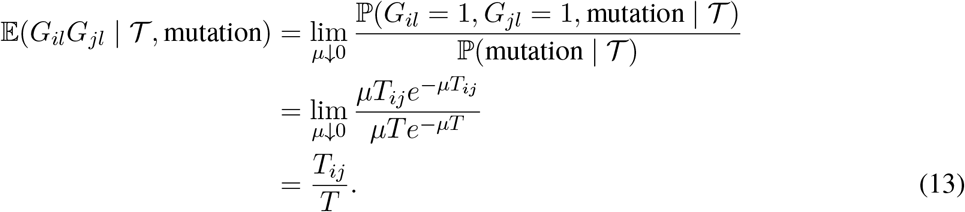

This fits the intuition that, conditional on a site being variable, individuals will share that mutation only if it occurs in a common ancestor of those two individuals. Finally, putting these pieces together

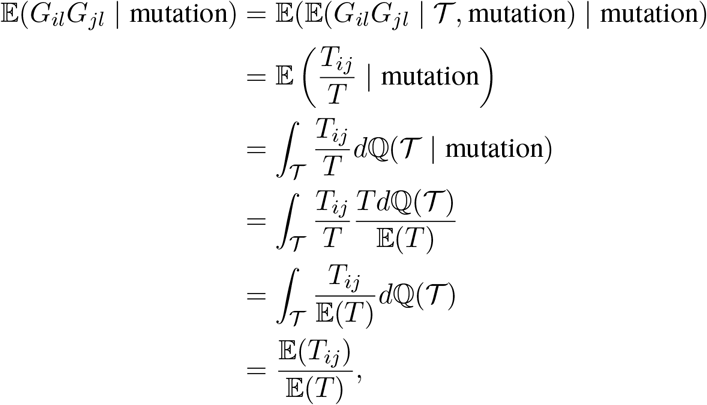

so that

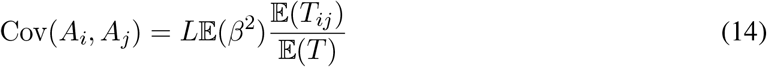

We note that this formula explicitly requires conditioning on the site being variable and that we are ignoring the effects of linkage disequilibrium among sites.

In principle, the entries of the relatedness matrix could be computed based on a demographic model; in this approach, one would be averaging over the randomness of both gene trees and mutations. This is the approach used by [91] to provide a genealogical interpretation of principal components analysis in genetics. However, demographic models do not typically capture fine-scale population structure. Instead, several recent methods in statistical genetics [58, 59, 93] and in phylogenetics [94] take as input a genome-wide inference of local gene trees. Then, the only source of randomness is averaging over the placement of mutations as in (13) and averaging over trees is accomplished by taking the average over observed gene trees. For example, Link and colleagues [58] use estimated gene trees in a region of the genome to compute the expectation of a local GRM formed from neutral variants falling on the estimated trees as a Poisson process. These matrices are then used as input to a variance-components model, which brings some advantages in mapping QTLs. Specifically, the resulting (conditional) expected genetic relatedness matrices naturally incorporate LD, providing better estimates of local genetic relatedness than could be formed from a handful of SNPs in a local region [58, 93].

### The phylogenetic variance-covariance matrix

In an extreme case, we might consider only variation among long-separated species. If we ignore incomplete lineage sorting, there may be only a single tree that describes the relationships among species, and the expectation over gene trees used in the previous subsection can be dropped, leaving us with (13). Then the entries of the relatedness matrix Σ, which in the case of phylogenetic methods is referred to as the phylogenetic variance-covariance (or vcv), are given by

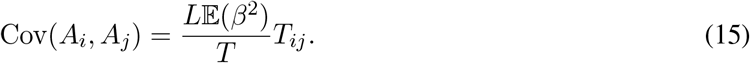

This can be recognized as the covariance under the Brownian motion model [45] commonly used to model continuous traits in phylogenetics, given a phylogenetic tree, when setting the diffusion rate *σ*^2^ of the Brownian motion process to

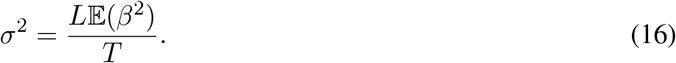

The formulation of equation 16 may look unfamiliar in phylogenetics, where the Brownian motion rate is typically taken to be *V*_*A*_*/N*, where *V*_*A*_ is the additive genetic variance and *N* is the effective population size, following Lande [95] or simply *U* 𝔼(*β*^2^), where *U* represents the total mutation rate toward causative alleles, following Lynch and Hill [96]. To reconcile our result with the existing literature, note that under the Poisson model described in the previous section 𝔼(*L*) = *UT*, so that

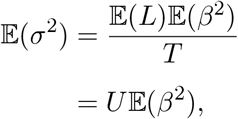

as shown by Lynch and Hill [96]. Further, under a neutral model, the equilibrium additive genetic variance *V*_*A*_ is proportional to *NU* 𝔼(*β*^2^) [97]. Thus, under neutrality,

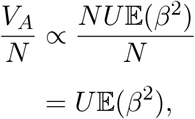

showing that under a neutral model, the Lande formulation is equivalent to the Lynch and Hill formulation, up to constants that depend on ploidy. Thus, we see that our equation 16 matches familiar formulations in the literature [98].

Consistent with previous arguments (e.g., [35]), this result also implies that one straightforward interpretation of the standard PGLS model is that it stratifies the regression between focal variables by an unobserved variable (or variables) that evolved primarily by drift. Hansen and colleagues have pointed out that this may not be an appropriate model for testing for adaptation [32, 33, 99], which was the primary motivation for developing many comparative methods in the first place [100]. Moreover, recently, standard PGLS has fallen into question in scenarios in which there is discordance between the gene tree and the species [101, 102]. Our formulation makes it clear that the standard PGLS formulation only applies when there is a single tree underlying all loci; if there is instead a distribution of gene trees, equation 14 suggests that the appropriate thing to do is to average over gene trees, as suggested by Hibbins *et al*. [102], and as done in a statistical genetics setting [58, 59]. Nonetheless, as we illustrate below, the fact that the standard phylogenetic regression falls out as a special case of the same general model as standard statistical genetics models is useful in practice, even if not always in theory.

### The connections among different approaches to modeling phenotypic covariance

Figure 1 provides a conceptual picture of how the various approaches are related to each other. The left side shows the situation typical in genome-wide association settings: SNP genotypes, shown as a matrix of variable sites with derived alleles colored in red, is determined by the topology of gene trees and the mutations that fall on them. The GRM is computed on the basis of the SNP genotypes, as in equation (8). If gene trees are known, then the eGRM can be computed by first averaging over Poisson placement of mutations as in equation (13) and subsequently taking an empirical average over gene trees. If only a demography is known, both gene trees and mutations can be averaged over using coalescent theory, as in equation (14). The right hand side shows the situation in phylogenetics: on a single fixed tree, the population trait mean evolves according to a Brownian motion. This results in a multivariate Gaussian distribution of phenotypes across species. We show that the covariance predicted by the Brownian motion model is equivalent to the covariance predicted by averaging over Poisson distributed mutations on a gene tree that is fixed to coincide with the species tree. In the figure, we highlight bifurcating population trees for simplicity and clarity, but the results also apply in complex demographic scenarios with admixture and reticulation.

**Figure 1:**
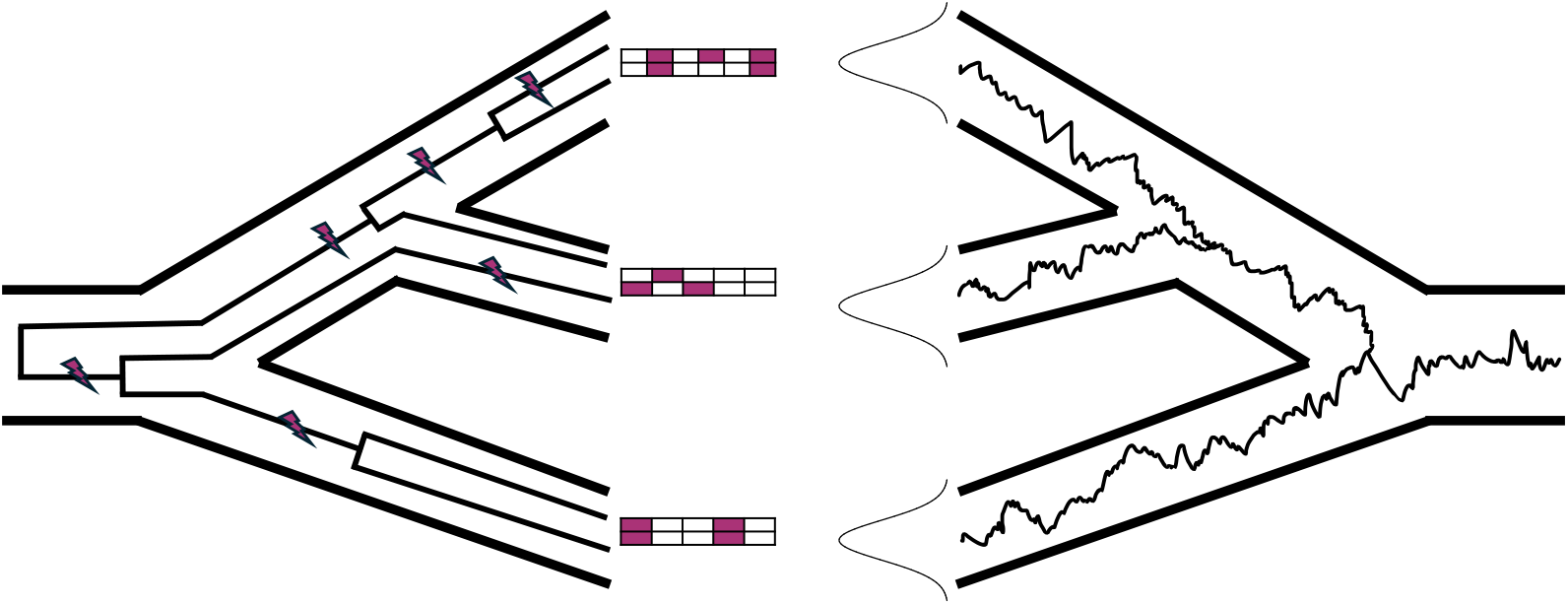
Relationship between different models of phenotypic covariance. The left-hand side shows the situation when multiple samples are taken from each group, as is the case in a genome-wide association study. The population tree is indicated by bold lines, and inside of it gene trees are indicated by thinner lines. Mutations on the gene trees are indicated by purple lightning bolts. The mutations on the gene tree result in genotype matrices, shown as one 2×5 array per species, with purple-filled entries indicating mutations. The right-hand side shows the situation in phylogenetics, where the species mean phenotype, indicated by a thin squiggly line, evolves according to a Brownian motion within a species tree, indicated by bold lines. The distribution of possible phenotypes within each species is marginally Gaussian.

### How the same type of unmodeled structure misleads both GWAS and phylogenetic regressions

The result of the previous section that standard models in statistical genetics and phylogenetics are closely related immediately suggests that these models might suffer the same pathologies under model misspecification, and that solutions to these pathologies could be shared across domains. Here we illustrate this by studying the problem of how unmodeled (phylo)genetic structure biases estimates of regression covariates. This problem has received much attention in both the statistical genetics [103, 104] and phylogenetics literature [34, 35, 105], but the approaches taken in the two fields differ.

We assume that we have a sample of size *n* with a vector of a predictor, *x* = (*x*_1_, *x*_2_, …, *x*_*n*_)^*T*^, and a trait, *y* = (*y*_1_, *y*_2_, …, *y*_*n*_)^*T*^. In the context of genome-wide association studies, *x* may be the (centered) genotypes at a locus to be tested for association, while in the context of phylogenetics, *x* is often an environmental variable or another trait that is hypothesized to influence *y*. Then, the regression model is

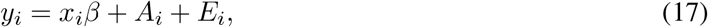

where *A*_*i*_ and *E*_*i*_ are the genetic and environmental components, as in equation 2, and *β* is the effect of *x* on *y*. Because of the correlation inherent in the genetic component, estimating the effect of *x* on *y* requires accounting for that covariance, or else hypothesis testing will not have the appropriate type I error rate [106, 107]. In genome-wide association studies, *β* is the effect size of the locus being examined, while in phylogenetics it may quantify the effect of an environmental variable or other continuous trait, rather than the effect of an allele.

### Theoretical analysis

To understand the purpose and limitations of corrections for (phylo)genetic structure, we examined the properties of the estimators of regression coefficients with and without correction for (phylo)genetic structure. The simplest estimator, 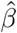, is the ordinary least squares estimator. In that case, the estimated effect size is

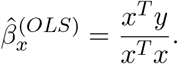

Because equation 17 shows that *y* is (phylo)genetically structured with covariance matrix Σ, we can expand 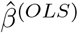 in terms of the eigenvectors and eigenvalues of Σ by diagonalizing,

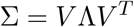

where *V* = [*v*_1_ *v*_2_ …*v*_*n*_] is a matrix whose columns are the eigenvectors of Σ, and Λ = diag(*λ*_1_, *λ*_2_, …, *λ*_*n*_) is a diagonal matrix whose diagonal is the eigenvalues of Σ. Σ, by virtue of being a covariance matrix, is guaranteed to be positive semidefinite. Thus, by the spectral theorem, the eigenvectors of Σ, will be able to form an orthonormal basis of ℝ^*n*^. In practice, Σ may have repeated eigenvalues, and hence the eigenvectors may need to be orthogonalized; intuitively, these repeated eigenvalues correspond to individuals, populations, or species that share the same evolutionary history. We proceed by assuming that the eigenvectors of Σ have been orthogonalized. Then,

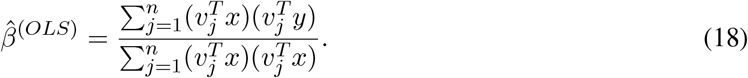

This shows that we can conceptualize the ordinary least squares estimator as adding up the correlations between *x* and *y* projected onto each eigenvector of Σ. Loosely, large-magnitude slope estimates arise when *x* and *y* both project with large magnitude onto one or more eigenvectors of Σ. If an eigenvector of Σ is correlated with a confounding variable, such as the underlying (phylo)genetic structure, then *x* and *y* may both have substantial projections onto it, even if *x* and *y* are only spuriously associated due to the confound.

Two seemingly distinct approaches have been proposed to address this issue. First, researchers have proposed including the eigenvectors of Σ as covariates. In the phylogenetic setting, this is known as phylogenetic eigenvector regression [108]. (In practice, researchers often use the eigenvectors of a distance matrix derived from the phylogenetic tree rather than Σ itself, but these two matrices have a straightforward mathematical connection [109]). In the statistical genetics setting, the analogous approach is to include the principal component projections of the data that are used to generate the genetic relatedness matrix—i.e., the principal components of the genotype matrix [23]—in the regression. To see that the eigenvectors of Σ are equivalent to the projections of the data onto the principal components, suppose we have *n* × *L* genotype matrix *G* whose rows represent individuals and whose columns represent genetic loci. Entry *i, j* contains the number of copies of a non-reference allele carried by individual *i* at locus *j*. Then, the projections of the data onto the *k*th principal component is given by

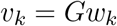

where *w*_*k*_ is the *k*th eigenvector of the *L* × *L* covariance matrix of the *genotypes*, i.e. *G*^*T*^ *Gw*_*k*_ = *λ*_*k*_*w*_*k*_. Then,

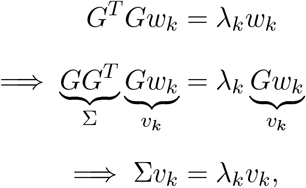

where the second line follows from left multiplying the first line by *G*. For clarity, we emphasize that this is distinct from computing eigenvectors of the phenotypic matrix (if multiple phenotypes are measured), which is intended to solve a different problem [110, 111]. However, if all traits evolve by identical Brownian motions, the eigenvectors from the phenotype matrix will coincide with the eigenvectors of Σ.

Now, we construct a design matrix *X* = [*x v*_1_ *v*_2_ …*v*_*J*_],whose columns are the predictor *x*, followed by the first *J* eigenvectors of Σ. (In theory, any set of eigenvectors of Σ might be included in the design matrix, but in statistical-genetic practice, it is typically the leading eigenvectors.) Then, using standard theory, the vector of coefficient estimates is

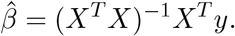

After some linear algebra (see Appendix), the estimate of the coefficient of the predictor *x* is

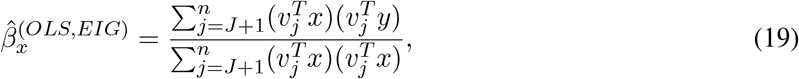

i.e., it is the OLS estimator, but the first *J* eigenvectors of Σ are removed. This shows why inclusion of the eigenvectors of Σ as covariates can correct for (phylo)genetic structure: it simply eliminates some of the dimensions on which *x* and *y* may covary spuriously. However, it also shows the limitations of including eigenvectors as covariates. First, because it is simply cutting out entire dimensions, it can result in a loss of power. Second, confounding that aligns with eigenvectors that are not included in the design matrix is not corrected.

The second approach to including the eigenvectors of Σ as covariates is to use Σ itself to model the residual correlation structure. In phylogenetic biology, this is accomplished using phylogenetic generalized least squares (PGLS) [39, 40], whereas in statistical genetics this is accomplished using linear mixed models (LMM) [28, 112]. (We work with generalized least squares below; for a similar argument in an LMM setting, see [27].) In both settings, it is common to add a “white noise” or “environmental noise” term, such that the residual covariance structure is 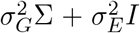, where 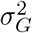 scales the contribution of genetics, 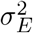 scales the contribution of environment, and *I* is the identity matrix. In the context of phylogenetics, the relative sizes of 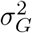 and 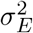 are of interest when estimating the phylogenetic signal measurement Pagel’s lambda [113, 114], whereas in statistical genetics, they are the subject of heritability estimation [115]. Then, both PGLS and LMM approaches model the data as

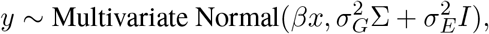

where 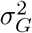 and 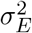 are typically estimated, for example by maximum likelihood [116], residual maximum likelihood [78], Haseman-Elston regression [117, 118], or other methods [28, 112, 119]; see Min *et al* for a comparison some estimation approaches and an examination of the impact of linkage disequilibrium [120].

For the theoretical analysis that follows, we assume 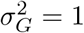 and 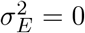. This does not restrict the applicability of our analysis, because 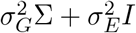 has the same eigenvectors as Σ, with corresponding eigenvalues 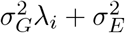, where *λ*_*i*_ are the eigenvalues of Σ.

With these assumptions, the regression coefficient can be estimated via generalized least squares,

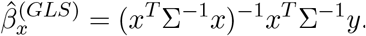

By diagonalizing Σ (see Appendix), the estimated regression coefficient is

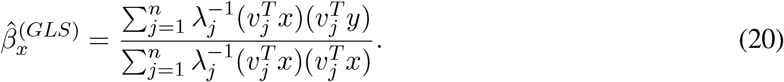

Like the ordinary least squares estimator in equation 18, this expression includes all the eigenvectors of Σ.

However, it downweights each eigenvector according to its eigenvalue. Thus, GLS downweights dimensions according to their importance in Σ, which aims to describe the structure according to which *x* and *y* may be spuriously correlated. However, unlike equation 19, it retains all dimensions. Compared with adjusting for the leading eigenvectors of Σ using OLS, the GLS approach retains some ability to detect contributions to associations that align with the leading eigenvectors. It also adjusts for Σ in its entirety, rather than just its leading eigenvectors. This means that it adjusts for even very recent (phylo)genetic structure, which will likely not be encoded by the leading eigenvectors. That said, one disadvantage of GLS is that it assumes that all eigenvectors of Σ contribute to confounding in proportion to their eigenvalues, potentially resulting in an inability to completely control for confounding if the effect of an eigenvector of Σ is not proportional to its eigenvalue, as may be the case with, for example, environmental confounding. In other words, the cost of including some adjustment for every eigenvector of Σ is an assumption as to how these eigenvectors relate to confounding. Other assumptions can be implemented, for example by transforming Σ, but model misspecification remains a risk once a specific model is chosen.

Where sample sizes and computational resources allow it, the state-of-the-art practice in statistical genetics is to use a linear mixed model framework while *also* including eigenvectors of Σ as covariates [28, 112, 119]. This at first may seem surprising, because it seems to be controlling for Σ twice. However, the analysis above suggests that including the eigenvectors as covariates and using generalized least squares have different, and perhaps complementary, outcomes. To see how they interact, again construct the design matrix *X* = (*x v*_1_ *v*_2_ … *v*_*J*_), whose columns are the predictor *x*, followed by the first *J* eigenvectors of Σ, and use the generalized least square estimator,

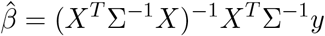

After some linear algebra (see Appendix), we again obtain the estimate of the regression coefficient of *x*,

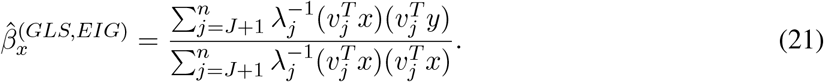

Thus, using the eigenvectors of Σ as covariates in a generalized least squares framework may provide the benefits of both approaches: if there is confounding in a eigenvector of Σ that is “too large”—that is, it is out of proportion with its associated eigenvalue—then if that eigenvector is included in the design matrix, it will simply be excised from the estimator, as in equation 19. However, we still maintain the ability to control for spurious association between *x* and *y* due to the structure of Σ but not along included eigenvectors, as in equation 21. The major difficulty is in identifying the eigenvectors of Σ that might be associated with confounding effects larger than their corresponding eigenvalues would suggest.

### Simulation analysis

To put the intution developed from the previous subsection into practice, we performed simulations in both phylogenetic and statistical-genetic settings. First, to explore how the different approaches outlined above correct for both (phylo)genetic structure and environmental confounding, we performed simulations inspired by Felsenstein’s “worst case” scenario [35, 46]. Felsenstein’s worst case supposes that there are two diverged groups of samples that are measured for two variables *x*, and *y*, which are then tested for association; the only (phylo)genetic structure is between the two groups. In the phylogenetic setting, we represent the two clades as star trees with 100 tips each, connected by internal branches, and we simulate *x* and *y* as arising from independent instances of Brownian motion along the tree (see Methods). In the statistical genetics setting, we use msprime [121] to simulate 100 diploid samples from each of two populations, and then simulated quantitative traits using the alpha model [22] (see Methods). In this setting, McVean [91] showed that the first eigenvector of Σ captures population membership; hence, we only include the first eigenvector to capture any residual confounding. To perform inference in the phylogenetic case, we used the package phylolm [116], and for the statistical-genetic case, we used a custom implementation of REML [78].

We first explored the impact of deepening the divergence between the two clades, starting from no divergence and increasing to high divergence (Figure 2A,C). As expected, we see ordinary least squares fails to control for the population stratification as the divergence time becomes large, resulting in high false positive rates. However, all of the other approaches appropriately controlled for the population stratification. This is as expected: in the case of two populations, all of the (phylo)genetic stratification is due to the accumulation of genetic variants in each group. Hence, either discarding the correlation between *x* and *y* on the dimension corresponding to group membership as in equation 19 or downweighting it as in equation 21 is sufficient to remove the confounding effect of the population stratification.

**Figure 2:**
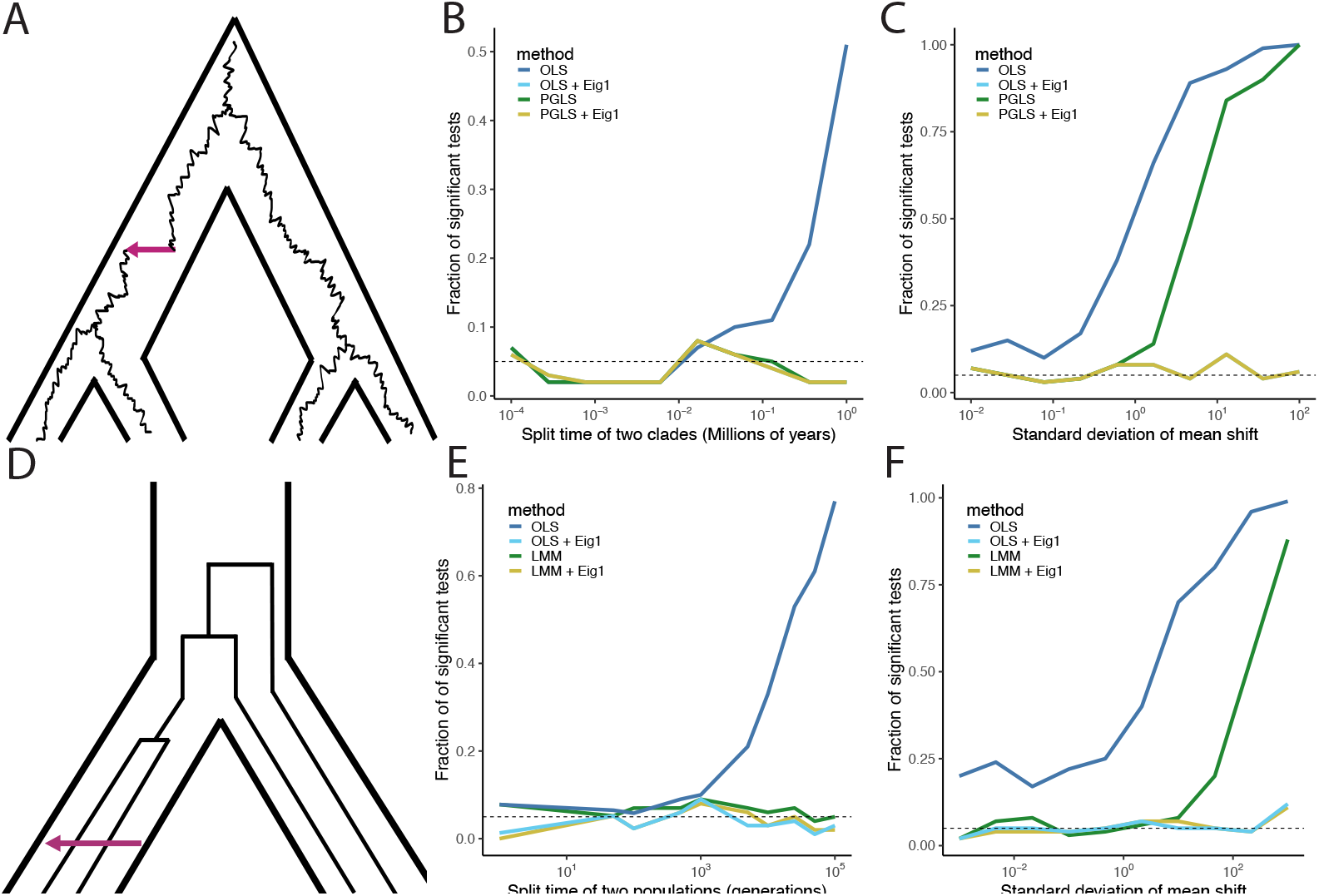
Performance of different methods for controlling confounding in Felsenstein’s worst case. A) A depiction of Felsenstein’s worst case in the phylogenetic setting. A Brownian motion evolves within a species tree separating two clades. For simplicity, two tips are shown in each clade; in the simulations, each clade contains 100 tips. The purple arrow shows a simulated singular evolutionary event (see text) B) The false-positive rate of each method in a simulated phylogenetic regression as a function of divergence time between the two groups. The horizontal axis shows the divergence time, while the vertical axis shows the fraction of tests that would be significant at the 0.05 level. Each line represents a different method. The lines for OLS + Eig1 and PGLS + Eig1 are completely overlapping. C) The false positive rate of each method in a simulated phylogenetic regression as a function of the size of non-Brownian shifts in both predictor and response variables. The horizontal axis shows the standard deviation of the normal distribution from which the shift was drawn, and the vertical axis shows the fraction of tests that would be significant at the 0.05 level. The lines for OLS + Eig1 and PGLS + Eig1 are completely overlapping. D) A depiction of Felsenstein’s worst case in the statistical genetic setting. Gene trees with mutations are embedded within a population tree depicting two divergent populations. For simplicity, two samples are shown within each population; in the simulations, each population consists of 100 diploid individuals. The purple arrow shows a simulated environmental effect (see text) E) The false-positive rate of each method in a simulated GWAS as a function of divergence time between the two groups. The horizontal axis shows the divergence time, while the vertical axis shows the fraction of tests that would be significant at the 0.05 level. Each line represents a different method. F) The false-positive rate of each method in a simulated GWAS as a function of the size of an environmental shift. The horizontal axis shows the standard deviation of the normal distribution from which the shift was drawn, and the vertical axis shows the fraction of tests that would be significant at the 0.05 level.

Despite the success of both OLS with eigenvector covariates and generalized least squares in controlling for population stratification, it has recently been recognized that phylogenetic generalized least squares does not control for all types of confounding in Felsenstein’s worst case: for example, if there is a large shift in *x* and *y* on the branch leading to one of the groups, phylogenetic generalized least squares produces high false positive rates [35]. Because including the first principal component will completely eliminate the contribution to the estimated coefficient that projects along group membership, whereas generalized least squares will only downweight it, we reasoned that including the first eigenvector in either ordinary or generalized least squares should restore control even in the presence of large shifts.

We tested our hypothesis using simulations with divergence time in which ordinary least squares was not sufficient to correct for population stratification. In the phylogenetic case, we simulated an additional shift in one of the clades for both *x* and *y* by sampling from independent normal distributions, while in the statistical-genetic case, we simulated an environmental shift sampled from a normal distribution in one of the clades (Figure 2B,D). As expected, ordinary least squares is insufficient to address the confounding, and becomes increasingly prone to false positives as the size of the shift increases. In line with our hypothesis, phylogenetic generalized least squares and linear mixed modeling also fail to control for the shift as it becomes large, while including just a single eigenvector in each case is sufficient to regain control over false positives.

The preceding analysis might suggest that including eigenvectors of Σ as covariates is sufficient to adjust for (phylo)genetic structure while also being superior to generalized least squares in dealing with environmental confounding. Recent work, however, suggests that inclusion of principal components may not be able to adjust for more subtle signatures of population structure [8, 15, 104, 122]. To explore this, we simulated both phylogenetic regression and a variant association test using a more complicated model of population structure. For the phylogenetic case, we simulated pure birth trees with 200 tips, while in the statistical genetics case, we simulated pure birth trees with 20 tips and sampled 10 diploids from each tip using `msprime`. Then, as before, we simulated using a Brownian motion model in the phylogenetic case, or an additive model for the statistical genetic case.

As expected, using ordinary least squares without any eigenvector covariates does not control for population structure in either the phylogenetic or the statistical-genetic setting, while the methods that use generalized least squares estimates of the regression coefficients appropriately model population structure (Figure 3). Although adding additional eigenvectors reduces the false positive rate of ordinary least squares, it does not reach the desired false positive rate of 5%. This is in line with our theoretical analysis: as seen in equation 19, including eigenvectors in ordinary least squares eliminates dimensions that explain the most genetic differentiation, but the correlations on the remaining dimensions are not adjusted. Because there is substantial fine-scale population structure in these simulations, removal of just a few dimensions with large eigenvalues is not sufficient to control for the subtle signature of population structure. In the phylogenetic setting, we expect that including additional eigenvectors would eventually gain control of false discoveries, but it may require including all of the eigenvectors and result in an overdetermined problem. On the other hand, in the population-genetic simulations, including additional eigenvectors will not increase control over false discoveries. There are two reasons for this. First, because Σ is estimated from the genetic data, the eigenvectors themselves are estimated. In practice, this means that eigenvectors corresponding to small eigenvalues are estimated poorly. Second, because we have 200 samples but only 20 populations, many of the samples share the same evolutionary history, and hence several eigenvectors share the same eigenvalue “in theory”—that is, if viewed from the perspective of the population tree rather than the realized gene trees or genotypes. Roughly speaking, in this simulation, there are only approximately 20 eigenvectors that correspond to “true” confounding. In practice, due to randomness of mutations and gene trees, these eigenvectors will not share identical eigenvalues, but will nonetheless correspond to genetic differentiation of individuals with shared evolutionary history, and hence are not able to correct for genetic confounding. This is reminiscent of the observation that in some human genetics datasets, only the first few eigenvectors stably capture genetic differentiation [123].

**Figure 3:**
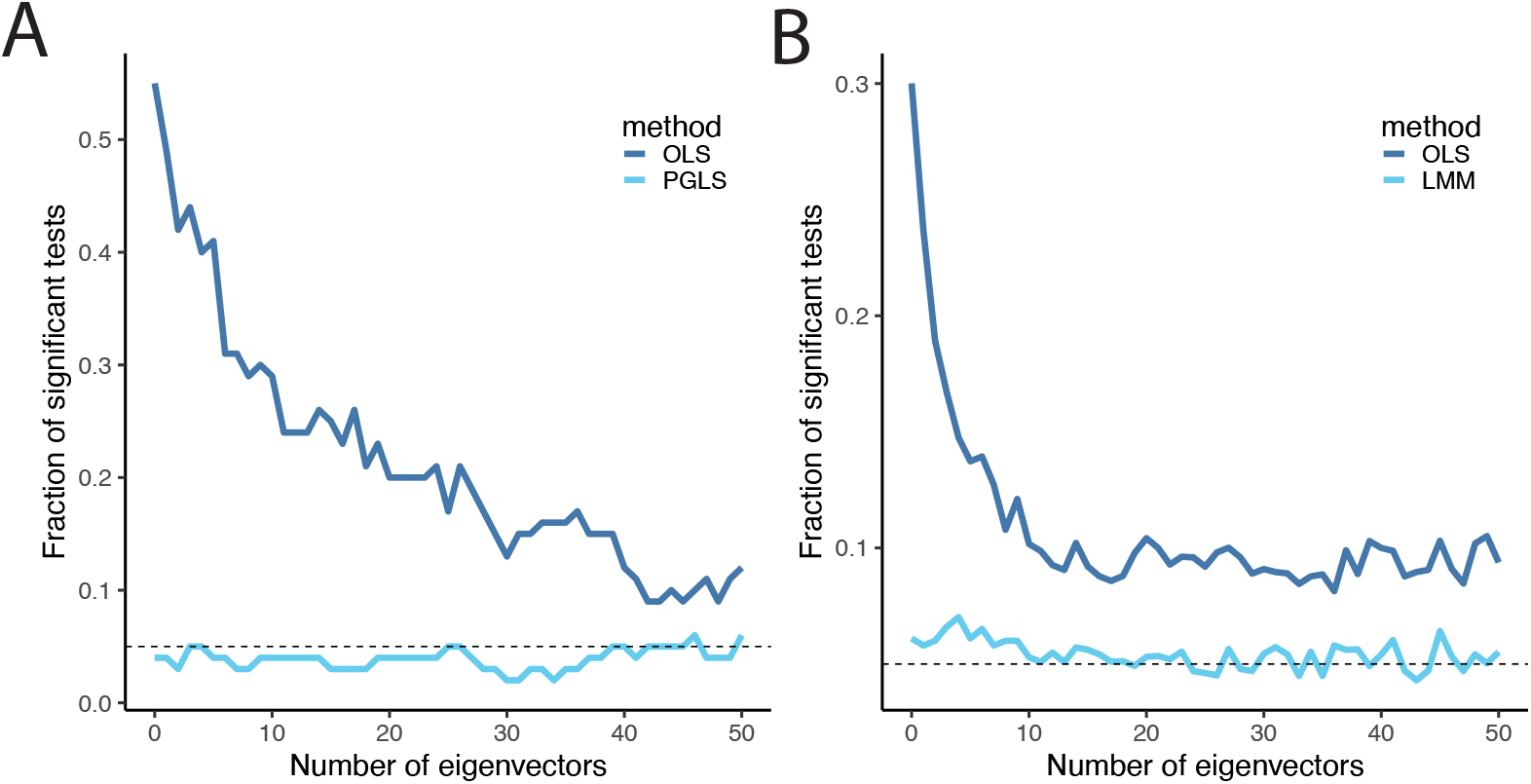
The performance of eigenvectors of the covariance matrix in a model with more complex population structure. A) Performance of ordinary least squares and phylogenetic least squares in a model with 200 tips related by a pure birth tree. The horizontal axis shows the number of eigenvectors included as covariates, and the vertical axis shows the fraction of tests that would be significant at the 0.05 level. B) Performance of ordinary least squares and a linear mixed model in a model with twenty populations related by a pure birth tree, and ten diploid individuals per population. The horizontal axis shows the number of eigenvectors included as covariates, and the vertical axis shows the fraction of tests that would be significant at the 0.05 level.

In contrast to including eigenvectors as fixed effects as part of an OLS analysis, generalized least-squares approaches, as shown in equation 21, will continue to correct for population structure that is found deeper into the eigenvectors of the correlation matrix (echoing points previously raised in the phylogenetics literature [124–126]). We also note that while the our analysis is focused on the eigenvectors of Σ, we suspect similar lines of reasoning may apply to other situations in which eigenvector regression is used, such as in spatial ecology [127].

### A case study of including eigenvectors as covariates in PGLS

Although the eigenvectors of the phylogenetic variance-covariance matrix (or closely related quantities) have often been included in regression models by researchers using phylogenetic eigenvector regression [108], to the best of our knowledge, phylogenetic biologists have not previously used these eigenvectors as fixed effects in a PGLS model—which we have shown above to be an effective strategy in theory. To illustrate the approach in practice, we re-examine a recent study by Cope *et al*. [128] that tested for co-evolution in mRNA expression counts across 18 fungal species. More specifically, these researchers were interested in testing whether genes whose protein products physically interacted (using independent data from [129]) were more likely to have correlated expression counts than those whose protein products did not. They found support for this prediction. While we suspect the core finding is robust, and there are some theoretical reasons to expect that RNA expression counts should be Brownian-like under some selective scenarios [130], other studies have shown expression counts for many genes in this dataset (and many others) are not well-described by a Brownian process [66, 131]. As such, some of their observed correlations could be spurious due to unmodeled phylogenetic structure [35].

We re-analyzed the data of Cope *et al*. [128] with the addition of the eigenvectors of (phylogenetic) Σ as fixed effects in the PGLS model (see Methods and Materials for details). Cope *et al*. used a correlated multivariate Brownian model to test their hypothesis, which is slightly different than the more common PGLS approach [132], but they are close enough for our purposes. We conducted several iterations of the analyses, varying the number of eigenvectors included from 1 to 10; Figure 4A shows how the different species project onto each principal component. We found that, as anticipated, the number of significant correlations decreased as more eigenvectors were included (Figure 4B). However, as more eigenvectors were included, the proportion of significant correlations in gene-expression count data in which the genes are known to physically interact increased (up to about 8 eigenvectors; Figure 4C). If we assume that the significant correlations for physically interacting genes are more likely to be true positives than those for pairs of genes not known to interact physically, then the results would suggest that including the eigenvectors in the analysis might reduce the false positive rate while still finding many of the true positives.

**Figure 4:**
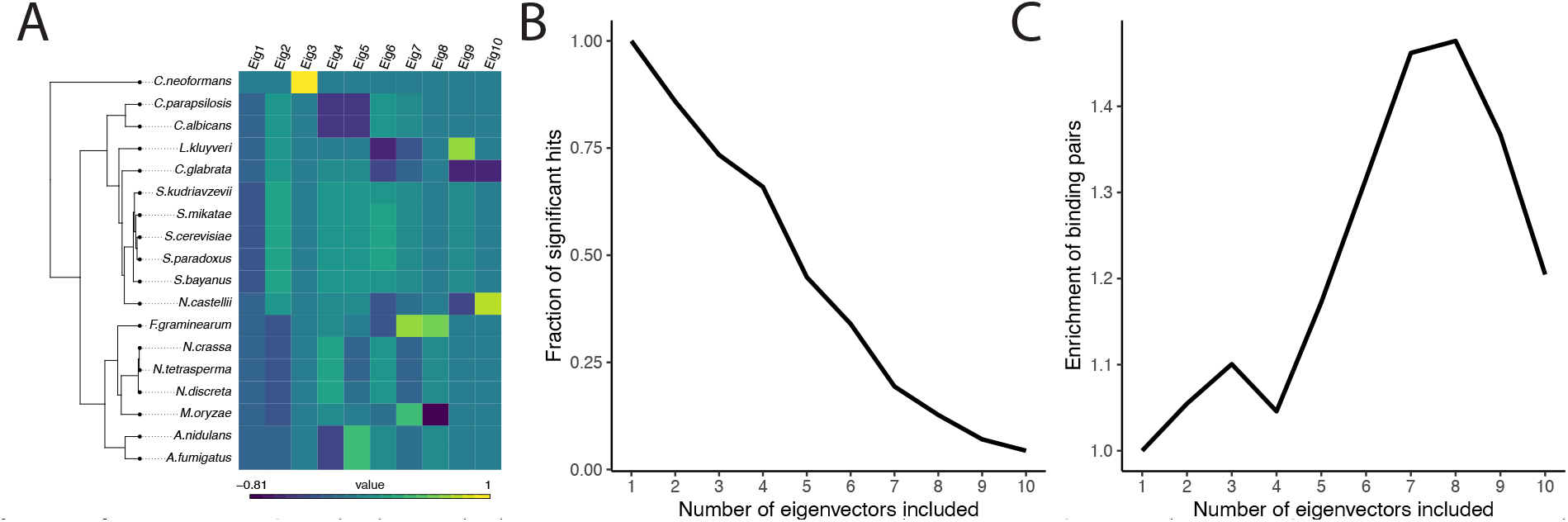
Impact of including phylogenetic eigenvectors on detection of coevolution of gene-expression levels in fungi. A) The fungal tree; colors indicate each species’ position in the first ten dimensions of principal component space. B) The overall number of significant pairs decreases as more eigenvectors are included in the regression. The horizontal axis indicates the number of eigenvectors included as fixed effects, and the vertical axis shows the proportion of significant pairs compared with a model that includes no eigenvectors as fixed effects. C) The enrichment of known binding pairs as a function of eigenvectors included. The horizontal axis indicates the number of eigenvectors included as fixed effects, and the vertical axis shows the enrichment of known binding pairs relative to a model in which no eigenvectors are included.

Uyeda *et al*. [35] suggest that one way to mitigate the spurious correlations arising from large, unreplicated events (see above sections) would be to simply use indicator variables in the regression model that encode the part of the phylogeny from which a tip descends (i.e., if there was a concordant shift in the means of both traits along a single branch, the indicator variable would be 0 if a species was on one side of that branch and 1 if on the other). This is similar in spirit to the use of hidden Markov models for the evolution of discrete traits [105, 133]. However, as Uyeda *et al*. point out, this leaves open the hard problem of identifying the branches on which to stratify. It is not possible to include an indicator for every branch, as the model would then be overdetermined. Using the simple method borrowed from GWAS studies of including eigenvectors of Σ as fixed effects in the typical phylogenetic regression may be a promising (partial) solution to the problem of spurious correlations.

## Discussion

### The genetic model vs. the statistical model

We began by adding assumptions to equation 2 in order to show that common practices in disparate areas of genetics can be seen as special cases of the same model. One notable assumption is that of a purely additive model [134] for the phenotype (equation 1). There are two reasons we might be suspicious of this assumption. First, it is debatable to what extent most traits obey the additive model, given evidence of non-additive genetic contributions to traits across species [135, 136]. However, even if non-additive contributions are important for determining individual phenotypes or for understanding traits’ biology, they might still contribute a relatively small fraction of trait variance, meaning they might be safely ignored for some purposes [137–139] (but see [140]). Second, we used a neutral coalescent model to find an expression for the Brownian motion diffusion parameter in terms of the effect sizes of individual loci (equation 16). Although this provides a satisfying justification for the use of a phylogenetic regression model with a Brownian covariance structure and for averaging over gene trees to accommodate ILS (*sensu* [102]), it is likely unreasonable in many situations. It has long been appreciated that, while a population-mean phenotype will be expected to evolve according to a Brownian process under simple quantitative-genetic models of genetic drift [43, 95, 98, 141] the Brownian rate estimated from phylogenetic comparative data is orders of magnitude too slow to be consistent with plausible values for the quantitative-genetic parameters used to derive the Brownian model [98, 141–143]. There are more elaborate explanations than pure genetic drift for why long-term evolution may show relatively simple dynamics [144] but understanding the coalescent patterns of loci under these scenarios is likely challenging [92] and beyond the scope of the present paper.

However, even if one finds the genetic model unreasonable, the equivalence of the *statistical* models used in statistical genetics and phylogenetics still holds: that is, the core structures of the models are the same, whether or not one is willing to interpret the parameters in the same way or not. Indeed, phylogenetic biologists have been here before, with the realization that PGLMMs are structurally equivalent to the pedigree-based analyses using the animal model from quantitative genetics [40–42] even though the recognition that they were equivalent did not rely on a specific genetic model for phenotypes (above, we prove that they can both be derived from the same genetic model). Nonetheless, the recognition of a structural equivalency between the animal model and the phylogenetic model made it possible to use techniques from quantitative genetics to solve hard problems in phylogenetic comparative methods. For example, inspired by a similar model from [145], Felsenstein developed a phylogenetic threshold model [146, 147], in which discrete phenotypes are determined by a continuous liability that itself evolves according to a Brownian process. Hadfield [148] proved this model was identical to a variant of the animal model and that existing MCMC algorithms could be used to efficiently estimate parameters and extend the threshold to the multivariate case, which had not been previously derived.

### Towards a more integrative study of the genetic bases of phenotypes

Building up this general framework is a step towards inference methods that coherently integrate intra- and interspecific variation to understand the genotype-to-phenotype map and how evolutionary processes, acting at different time scales, shape it. Indeed, the importance of evolutionary conservation in triaging functional variants in the human genome has long been appreciated and is becoming increasingly important as we collect larger samples of people; the same is true for the use of genomics in agriculture [57] and conservation genetics [55]. Recent work showed that evolutionary conservation accounts for the vast majority of the predictive power of a state-of-the-art deep learning approach to variant annotation [149, 150]. But most of the cutting-edge phylogenomic approaches for triaging variants typically do not use the phylogeny at all (i.e., only multiple sequence alignments [MSAs] are used), or include the phylogeny without an explicit evolutionary model [151]. This is a limitation because we are not making the most of the information in the tree, nor are we able to draw specific inferences about how evolutionary processes have shaped complex traits from the MSA alone. Overcoming this limitation is not straightforward and will require mechanistic modeling: the observed level of conservation is a nonlinear function of the strength of selection acting against variants at a locus; small changes in the strength of negative selection can greatly decrease the amount of variability seen on phylogenetic timescales, and this can cause counter-intuitive behavior of conservation scores [54, 152].

A key difficulty in combining information across timescales arises from different assumptions about evolutionary process. For example, the canonical GRM in statistical genetics assumes that the variance of an allele’s effect size is inversely proportional to the heterozygosity at the locus. As we show in the Appendix, this assumption can be justified under a model of mutation-selection balance with Gaussian stabilizing selection. However, we do not generally understand how robust such approaches are under more complex (and realistic) evolutionary scenarios that include the influence of genetic drift, nor how errors influence down-stream inferences [75, 79, 153, 154]. There is substantial evidence that rarer variants tend to have larger effect sizes [79, 155–159], which is broadly consistent with the motivation for the canonical GRM and for the more general “alpha model,” which supposes that the variance of the effect size of an allele is given as a power law function of its heterozygosity [22, 74, 78, 79]. However, close examination of GWAS effect sizes suggests a poor fit of the alpha model for many traits [153]. Further, recent explosive human population growth has resulted in a massive number of rare variants that have not been filtered by natural selection [160– 164]—many of these variants are rare not because they have been driven to low frequency by selection, but simply because they are the signature of mutation-drift balance in a growing population. As such, using the alpha model may result in over-estimation of heritability for traits where there is a substantial contribution of genetic drift and may result in incompletely controlled confounding in trait mapping studies. And although effect sizes of individual causal variants can be estimated well for common variants, this is unlikely to ever be possible for rare variants; hence, a realistic model of effect sizes as a function of allele frequency is necessary for inclusion in efforts such as rare-variant association studies [52, 165–167].

In contrast, in our derivation of gene-tree (i.e., those using the eGRM) and phylogenetic (i.e., using the phylogenetic vcv) model, we assume that effect sizes and genotypes were uncoupled, and that mutations fall on a gene tree as a Poisson process [91]. These assumptions imply that the causal variants are neutral. This is both inconsistent with the assumptions used in the above-mentioned case of the GRM, and contradicts a wealth of evidence from both within and among species that quantitative trait variation is under some form of selection [168–175] and that the effect sizes of causal variants tend to be larger in more evolutionarily conserved regions [50, 149, 176–179], which also implies an important role of purifying selection. The alpha model, or presumably other models of the relationship between effect size and allele frequency, can be incorporated in an eGRM [59, 93]. After all, an eGRM is an expectation (under Poisson-process mutation) of a GRM, and so any scaling applied to genotypes in computing the GRM can be made to apply to the eGRM. But the interpretation becomes complicated: Poisson mutations only occur under neutrality, and the effect sizes of mutations seem to “look into the future” to determine how many tips are below the mutation. Theoretical work is needed to understand the implications of incorporating the alpha model in eGRMs. One way phylogenetic biologists include selection is by modeling the evolution of quantitative traits with an Ornstein-Uhlenbeck (OU) process [99, 180–183], which can be derived from a quantitative-genetic model of stabilizing selection [95], although in practice, the OU model is often interpreted more of a phenomenological model of the evolution of the adaptive peaks [44, 184] than as literally representing stabilizing selection. Under an OU model, phenotypes on a phylogeny will be drawn toward a common optimum value; hence, covariances in Σ are higher than expected under a Brownian motion model. Many researchers have used the Σ matrix derived from an OU process in PGLS models [180, 185]; this is straightforward, because the data remains multivariate Gaussian [39, 116]. One could potentially use an analogous approach to model phenotypic evolution along gene trees within a species (to inform the construction of eGRM, for example). This would improve inferences from both tree-based GWAS (sensu [58, 59]) and from emerging phylogenetic comparative approaches that consider gene trees rather than just the species trees [101, 102, 186] (such approaches are important as only using a single species tree may lead one to mistake similarity due to common ancestry for convergence [63, 187, 188]).

We suspect that there are additional connections between statistical genetics and phylogenetics that we have not mapped out here and that could be profitably explored. For example, in most of the applications in which phylogenomic data are used to inform mapping studies, researchers have large-scale phenotypic and genomic sampling for a focal population or species and then sparser genomic sampling (often a single genome) and an estimate of phenotypic means (if even that) for the others. However, there are emerging datasets from closely related species that have dense phenotypic and genomic samples from multiple lineages [189, 190]. We anticipate that our framework could be used to derive more principled and powerful approaches for analyzing these types of data. At the other extreme are methods in which we have sparse sampling of both phenotypes and genomes for a phylogenetically diverse set of species (which generally fall under the PhyloG2P label, mentioned above [60]. In this case, researchers either use phylogenetic data to uncover convergent mutations associated with phenotypic convergence across lineages (e.g., [191]) or more commonly, identify regions with a relatively large number of substitutions—but not necessarily the same ones—in phylogenetically distinct lineages that have convergently evolved the same phenotype [192, 193]. For example, Sackton and colleagues [194] used such an approach to identify regulatory regions that had high rates of evolution in lineages of flightless birds; they also demonstrated that some of these regions influence wing development using experimental perturbations. Such rate association tests (see also [195]) seem to be very similar, both conceptually and statistically, to techniques used in rare-variant association studies, which look for local enrichment of rare variants in cases vs. controls, rather than associating single variants with phenotype [52, 165–167]. We suspect that one could derive a formal equivalence between these sets of methods as we did between GWAS and PGLS above using similar techniques.

There are clear biological rationales explaining why various types of analyses will be more or less informative at different timescales. But this is a difference of degree and not of kind. And the different methodological traditions in statistical genetics and phylogenetics are just that—traditions. There is no reason a researcher should think about the problem of trait mapping in a fundamentally distinct way just because she happened to be trained in a statistical genetics or phylogenetics lab. Ultimately, we should work to take the best ideas from both of these domains and blend them into a more cohesive paradigm that will facilitate richer insights into the molecular basis of phenotypes.

## Materials and Methods

### Simulation details

To perform phylogenetic simulations, we used the `fastBM` function from the `phytools R` package [196]. In all cases, Brownian motions were simulated independently and with rate 1. When performing phylogenetic simulations of Felsenstein’s Worst Case, we used `stree` from `ape` [197] to simulate two star trees of 100 tips, where each tip in the star tree had length 0.5. We then connected the two star trees using internal branches of varying length. To add a non-Brownian confounder, in each simulation we added an independent normal random variable with varying standard deviations to the *x* and *y* values for individuals from clade 1. (Within a given simulation, all individuals in clade 1 were augmented by the *same* value for each trait, while between simulations, the confounding effect was a random draw.) When performing simulations in a more complicated phylogeny, we used `TreeSim` [198] to generate pure-birth trees with birth rate = 1 and complete taxon sampling. Each simulation replicate used a different tree. For ordinary least squares on phylogenetic data, we used the `R` function `lm`. For PGLS on phylogenetic data, we used the `R` package `phylolm` [116] with the Brownian motion model and no environmental noise.

To perform genome-wide association study simulations, we first generated neutral tree sequences and mutations using `msprime` [121]. To ensure our results were not simply due to genetic linkage, we simulated a high recombination of 10^−5^ per generation with a mutation rate an order of magnitude lower, 10^−6^ per generation. We first simulated causal variants on a sequence of length 100000, and generated phenotypes by sampling an effect size for each variant from a normal distribution with mean 0 and variance 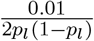 where *p*_*l*_ is the allele frequency of variant *l*. We then created each individual’s phenotype using the additive model, equation 1. We then added environmental noise so that the trait’s heritability was less than 1. In all simulations, every population had diploid population size 10000. To simulate the variant being tested for association, we simulated independent tree sequences and mutations and selected a random variant with allele frequency greater than 0.1. When simulating a GWAS analogue of Felsenstein’s Worst Case, we drew 100 diploid samples from each population, and varied the divergence time of the two populations. To include an environmental shift in one population, we added a normal random variable with varying standard deviation only to individuals in population 1. To simulate under a more complicated population structure, we simulated 20-tip pure birth trees using `TreeSim` with a birth rate of 5. We then multiplied all branch lengths by 10000 to convert them into generations, and imported them into `msprime` using the `from_species_tree` function. We then generated tree sequences and mutations, sampling 10 diploid individuals from each population. Note that each replicate simulation was performed on an independent random population tree. We performed association testing using a custom `python` implementation of the linear mixed model. We first used restricted maximum likelihood to estimate 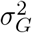 and 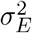, followed by using generalized least squares to estimate the regression coefficients and their standard errors.

### Phylogenetic analysis of yeast gene expression data

We obtained the species tree, gene expression matrix, and list of physically interacting genes from https://github.com/acope3/GeneExpression_coevolution [128]. We then randomly subsampled 500 genes that had measurements in at least 15 of the 20 species to test for association, resulting in 124750 pairs. Because of differential missingness among genes, we computed phylogenetic eigenvector loadings only on the subtree for which both genes had data present, meaning that each pair may have had slightly different eigenvector loadings. We then used `phylolm` [116] with no measurement error to estimate the regression coefficient. For each number of eigenvectors included, we corrected for multiple testing by controlling the FDR at 0.05 using the Benjamini-Hochberg procedure [199].

## Data availability

Code used to generate the results in this paper can be found at https://github.com/Schraiber/PGLS_GWAS

## Acknowledgements

We thank Emily Josephs, Alvina Adimoelja, Matt Aguirre, Garyk Brixi, Nikhil Milind, Roshni Patel, Julie Zhu, Katalin Voss, Graham Coop, and members of the Coop lab for comments on the manuscript, and Matt Hahn, Nick Mancuso, Jeff Spence, Sasha Gusev, Loren Rieseberg, and members of the Pennell, Edge, and Mooney labs for their thoughtful comments on parts of this study. Alex Cope provided additional guidance on our analysis of the yeast gene expression data. We acknowledge support from NIH grant R35GM137758 to MDE and R35GM151348 to MP.

## Appendix

### The relationship between effect size and allele frequency under strong Gaussian stabilizing selection

The derivation found here follows some of the development leading to equation 14a of Bulmer [200]. Under Gaussian stabilizing selection, the the fitness of an individual with phenotype *y* is given by

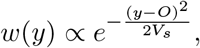

where *O* is the optimal phenotype and *V*_*s*_ is the variance of the stabilizing selection kernel (stronger selection is indicated by smaller *V*_*s*_). Assuming the population mean is near the optimum, an allele with effect size *β* will evolve according to underdominant dynamics, in which the minor allele is disfavored with with selection coefficient 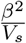, indicating that alleles that cause larger deviations from the optimum are disfavored, and the strength of selection on individual alleles is related to the strength of selection on the phenotype as a whole [77, 201, 202]. Mutation between alleles occurs (symmetrically) at rate *μ*. Thus,

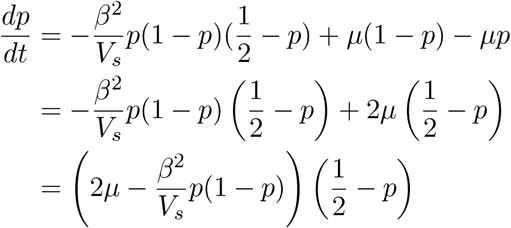

To find an equilibrium, set 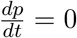. Thus, either

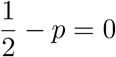

or

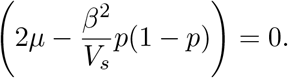

The first equation corresponds to an unstable equilibrium. Solving the second equation for *β*^2^ yields

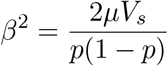

as desired.

### Covariance of the genetic component of the phenotype with pedigree-based kinship

With a known pedigree but no genotype data, we might derive the covariances among individuals in the genetic component of a trait by fixing the (unobserved) effect sizes and allele frequencies. For simplicity, we will also assume that individuals are not inbred. Given the pedigree, at any given locus, diploid individuals *i* and *j* have some probability of inheriting 0, 1, or 2 alleles identical by descent (IBD). Call these probabilities *r*_0_, *r*_1_, and *r*_2_, respectively. For example, for a parent-offspring pair, *r*_1_ = 1, and for a pair of full siblings, *r*_0_ = 1*/*4, *r*_1_ = 1*/*2, and *r*_2_ = 1*/*4.

Following equation 4, the covariance of the genetic component of the trait for individuals *i* and *j* is

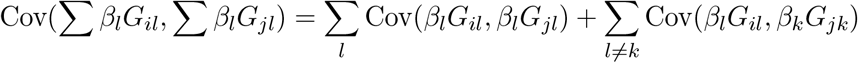

As in the main text, we assume that genotypes at distinct loci are independent, causing the second term to vanish. With the effect sizes treated as fixed constants, we have

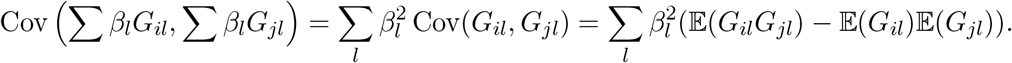

The *G* values are allelic counts of a non-reference allele (we assume the locus is biallelic), and if we fix the non-reference allele frequencies *p*_*l*_, then 𝔼(*G*_*il*_) = 𝔼(*G*_*jl*_) = 2*p*_*l*_

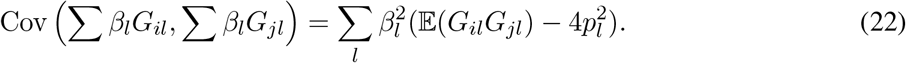

To proceed, we need 𝔼(*G*_*il*_*G*_*jl*_) given the IBD probabilities *r*_0_, *r*_1_, and *r*_2_. We thus compute the desired expectation conditional on each IBD state. *G*_*il*_*G*_*jl*_ = 1 if individuals are heterozygous, *G*_*il*_*G*_*jl*_ = 2 if one individual is heterozygous and the other is homozygous for the non-reference allele, and *G*_*il*_*Gjl* = 4 if both individuals are homozygous for the non-reference allele. If individuals *i* and *j* share no alleles IBD, using Hardy–Weinberg genotype probabilities (per the assumption of no inbreeding), the conditional expectation is therefore

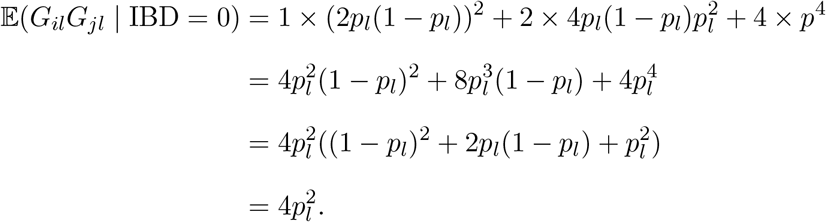

A similar calculation given one allele inherited IBD gives

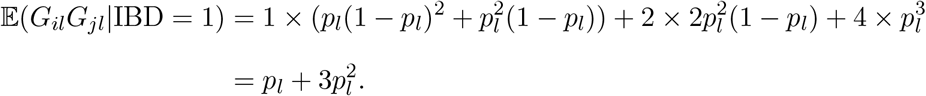

(To explain the first line: for the first term, the individuals are both heterozygous if the IBD allele is of the non-reference type and the other two are reference alleles, or if the IBD allele is non-reference and both of the non-IBD alleles are reference alleles. For the second term, since one individual is homozygous for the non-reference allele, the IBD allele must be of the non-reference allele, and of the other two alleles, one must be of each type. For the third term, both individuals are homozygous if the IBD allele and both non-IBD alleles are non-reference alleles.)

If the individuals share two alleles IBD, then

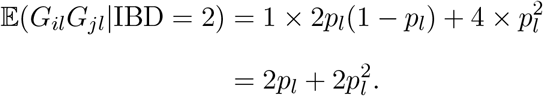

Combining these terms weighted by the IBD probabilities gives the desired expectation,

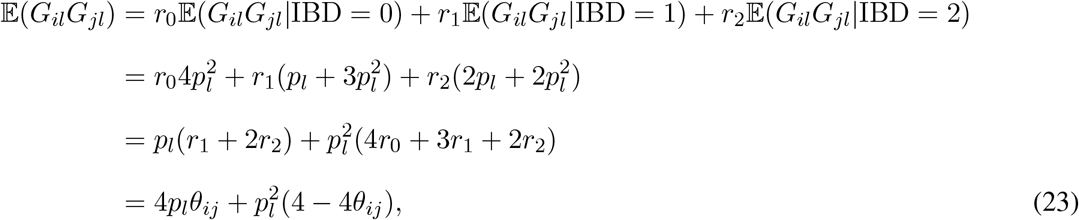

where the last line comes from noticing that *r*_0_ + *r*_1_ + *r*_2_ = 1 and defining the kinship coefficient *θ*_*ij*_ = *r*_1_*/*4 + *r*_2_*/*2, equal to the probability that a pair of alleles chosen at random, one from individual *i* and one from individual *j*, is IBD.

We can now return to the main covariance of interest by plugging the expression for 𝔼(*G*_*il*_*G*_*jl*_) from equation 23 into equation 22, giving

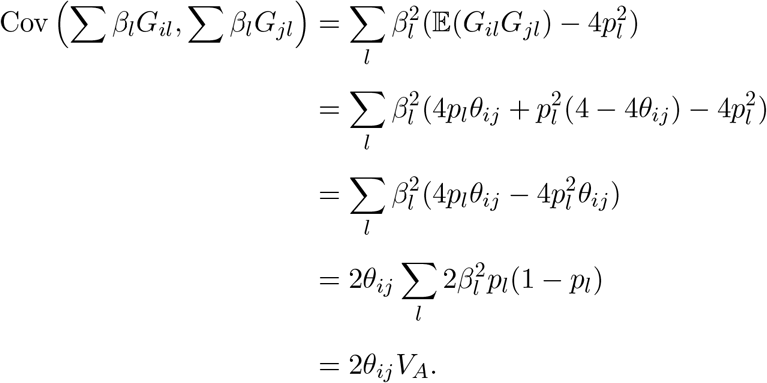

The final line comes from noting that the sum in the previous line is the additive genetic variance (*V*_*A*_) under the assumptions here, that is, the variance of the genetic component of the phenotype among outbred individuals. This is the result required in the main text.

### Covariance of genotypes when effect sizes and allele frequencies are independent

We have

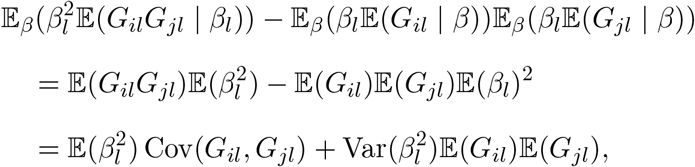

where the second line comes from the assumption that genotypes and effect sizes are independent, and the third line comes from adding and subtracting 𝔼(*G*_*il*_)𝔼(*G*_*jl*_)𝔼(*β*^2^) to the second line and simplifying. Further making the assumption that 𝔼(*β*) = 0,

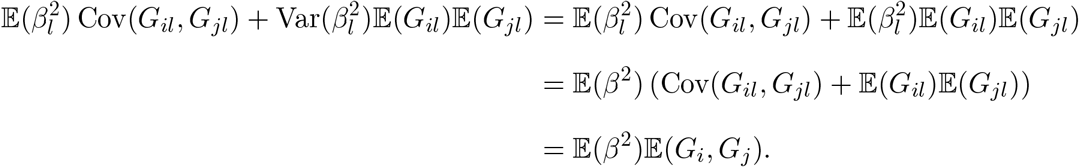

### The estimated regression coefficient from ordinary least squares with eigenvectors of the covariance matrix as fixed effects

Recall the design matrix *X* = [*x v*_1_ *v*_2_ …*v*_*J*_],where the *v*_*i*_ are orthonomal eigenvectors of Σ. Then,

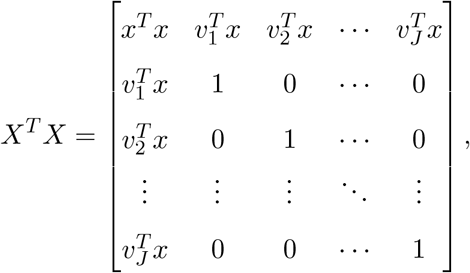

i.e. the first row and columns are the projection of *x* along each eigenvector and the rest of the matrix is an identity matrix. The identity matrix arises because the eigenvectors annihilate each other. Then,

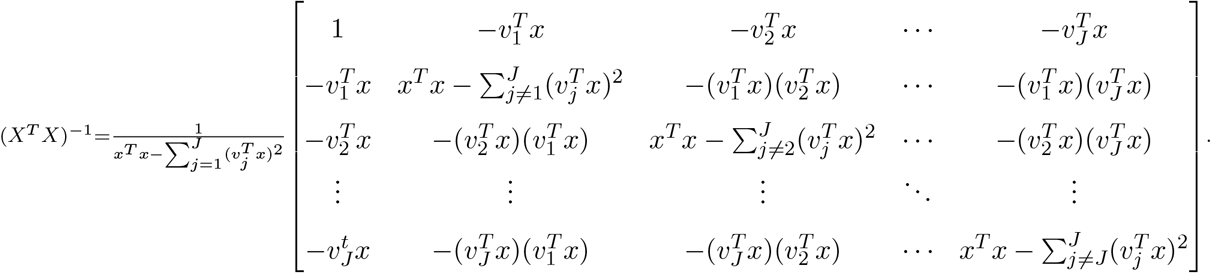

Ultimately, we only need the first row of (*X*^*T*^ *X*)^−1^ to estimate the regression coefficient for *x*. Note that by expanding *x*^*T*^ *x* in terms of the eigenvectors, we have that

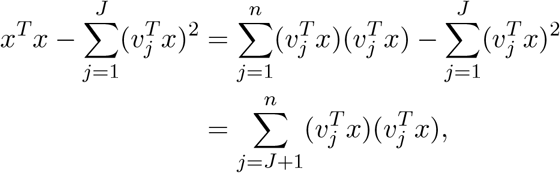

so that the first row of (*X*^*T*^ *X*)^−1^ is

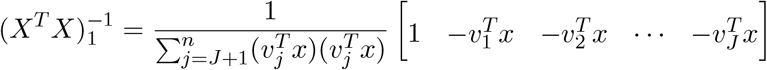

We also compute

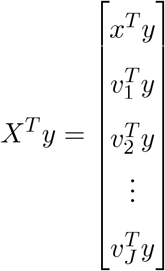

so that

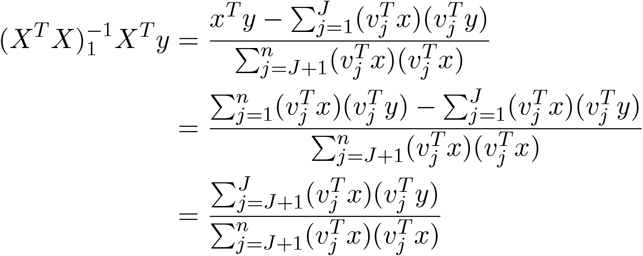

as desired. Note that when *J* = 0, this recovers the ordinary least squares estimator (equation 18), as expected.

### The estimated regression coefficient from generalized least squares

Recall that we can diagonalize the covariance matrix Σ as

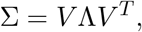

where *V* =[*v*_1_ *v*_2_ …*v*_*n*_] is a matrix whose columns are the eigenvectors of Σ and Λ = diag(*λ*_1_, *λ*_2_, …, *λ*_*n*_) is a diagonal matrix whose entries are the eigenvalues of Σ. If we have design matrix *X* =[*x v*_1_ *v*_2_ …*v*_*J*_], we can rewrite the genrealized least square estimator as

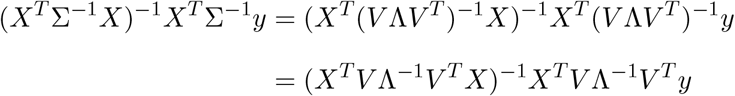

where 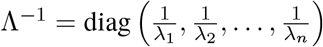, using the fact that *V* is an orthonormal basis. Next, compute

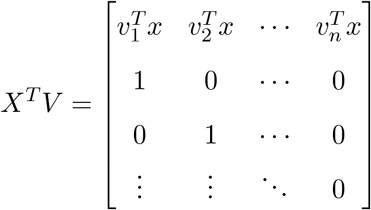

which is an (*J* + 1) × *n* matrix, so the pattern extends *J* + 1 rows and the identity elements come from the eigenvectors annihilating each other. Then,

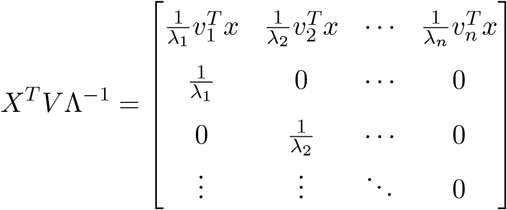

Note that *V* ^*T*^ *X* = (*X*^*T*^ *V*)^*T*^, so that

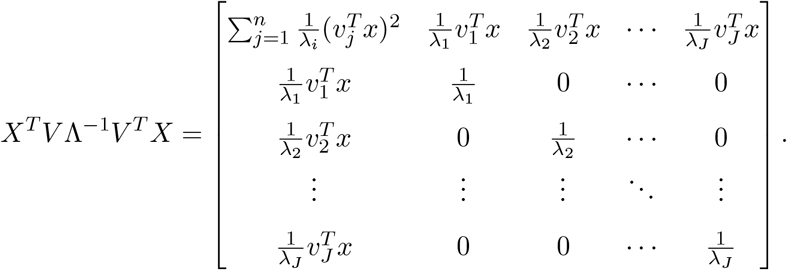

As in the case for ordinary least squares, we will only need the first row of the inverse,

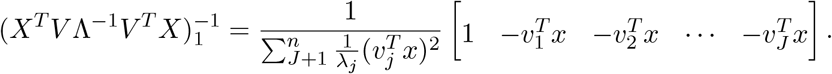

Next, compute

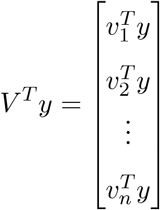

so that

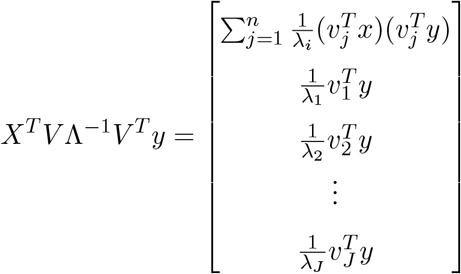

and finally

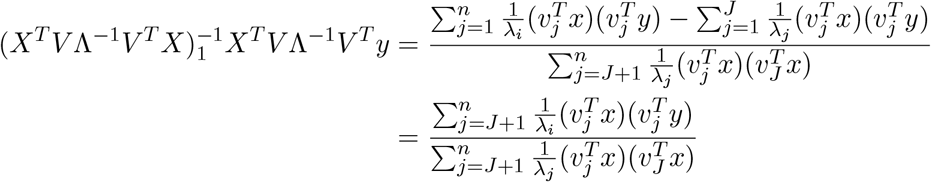

as desired. Setting *J* = 0 recovers the GLS estimator without any eigenvectors included as covariates, as in equation 20.

